# Monocyte-lymphocyte cross-communication via soluble CD163 directly links innate immune system activation and adaptive immune system suppression following ischemic stroke

**DOI:** 10.1101/144063

**Authors:** Grant C. O’Connell, Connie S. Tennant, Noelle Lucke-Wold, Yasser Kabbani, Abdul R. Tarabishy, Paul D. Chantler, Taura L. Barr

## Abstract

CD163 is a scavenger receptor expressed on innate immune cell populations which can be shed from the plasma membrane via the metalloprotease ADAM17 to generate a soluble peptide with lympho-inhibitory properties. The purpose of this study was to investigate CD163 as a possible effector of stroke-induced adaptive immune system suppression. Liquid biopsies were collected from ischemic stroke patients (n=39), neurologically asymptomatic controls (n=20), and stroke mimics (n=20) within 24 hours of symptom onset. Peripheral blood ADAM17 activity and soluble CD163 levels were elevated in stroke patients relative to non-stroke control groups, and negatively associated with post-stroke lymphocyte counts. Subsequent *in vitro* experiments suggested that this stroke-induced elevation in circulating soluble CD163 likely originates from activated monocytic cells, as serum from stroke patients stimulated ADAM17-dependant CD163 shedding from healthy donor-derived monocytes. Additional *in vitro* experiments demonstrated that stroke-induced elevations in circulating soluble CD163 can elicit direct suppressive effects on the adaptive immune system, as serum from stroke patients inhibited the proliferation of healthy donor-derived lymphocytes, an effect which was attenuated following serum CD163 depletion. Collectively, these observations provide novel evidence that the innate immune system employs protective mechanisms aimed at mitigating the risk of post-stroke autoimmune complications driven by adaptive immune system overactivation, and that CD163 is key mediator of this phenomenon.

## Introduction

Stroke triggers a systemic inflammatory response which results in a dramatic shift in the phenotype of the peripheral immune system. The innate arm of the peripheral immune system undergoes rapid activation, a phenomenon which results in significant elevations in circulating neutrophil and monocyte counts in the hours following onset.^1,2^ Conversely, the adaptive arm of the peripheral immune system shifts into a state of suppression, often characterized by a prolonged period of lymphopenia and limited antigen responsiveness.^2–4^ While the pathophysiological role of the innate immune response to stroke has been long appreciated, a growing body of evidence suggests that the adaptive immune response to stroke also critically influences patient outcome.

It is likely that the state of adaptive immune suppression which develops in response to stroke serves to limit the possibility of an autoimmune response as the blood brain barrier becomes disrupted and peripheral populations of lymphoid cells become exposed to unfamiliar central nervous system (CNS) peptides via the lesional and lymphatic accumulation of activated innate antigen presenting cells loaded with neural antigens.^5,6^ Expectedly, heightened adaptive immune activity immediately following stoke has been shown to prolong the inflammatory state and promote secondary tissue damage, factors which can ultimately contribute to poor prognosis.^6,7^ While the state of adaptive immune suppression is acutely protective, if prolonged, it can constitute a significant detriment to recovery.^8–12^ Patients which display prolonged adaptive immune suppression following stroke become highly susceptible to post-stroke infection,^13–16^ which is one of the leading causes of death in the post-acute phase of care.^9^ Despite the fact that it is becoming increasingly evident that this delicate balancing act between maintenance of self-tolerance and pathogen susceptibility plays a significant role in recovery, the mechanisms which drive the peripheral adaptive immune response to stroke are not yet fully understood.

CD163 is a membrane-bound scavenger receptor for extracellular hemoglobin which is believed to be expressed exclusively within the innate immune system,^17^ where it is predominantly found on monocytes and macrophages,^18^ and to a lesser extent on neutrophils.^19^ Various stimuli can trigger CD163 ectodomain shedding via cleavage by the metalloprotease ADAM metallopeptidase domain 17 (ADAM17),^20,21^ resulting in generation of a soluble truncated peptide (sCD163) which has been shown in multiple studies to directly interact with lymphocytes and inhibit antigen-induced proliferation.^22–24^ Interestingly, genome-wide transcriptomic screening performed by our group recently identified CD163 as being robustly up-regulated in the peripheral blood of ischemic stroke patients within hours of system onset.^25^ Furthermore, other groups have reported heightened ADAM17 activity in animal models of ischemic brain injury,^26–28^ as well as stroke-induced increases in peripherally circulating levels of other ADAM17 substrates such as tumor necrosis factor alpha (TNFα) in human subjects.^29,30^ Therefore, it is possible that stroke induces a rise in peripherally circulating levels of sCD163 via coordinate increases in CD163 expression and ADAM17 activity; such a rise in sCD163 levels could subsequently contribute to suppression of the adaptive immune system via the inhibitory effects of sCD163 on lymphocyte activity. Thus, the primary objective of this study was to investigate sCD163 as a potential modulator of post-stroke adaptive immune suppression.

## Methods and results

### Clinical and demographic characteristics

A cohort of 39 ischemic stroke patients was recruited along with two control groups, the first consisting of 20 neurologically asymptomatic subjects, and the second consisting of 20 acute stroke mimics (Table 1). Ischemic stroke patients were significantly older than both neurologically asymptomatic controls and stroke mimics. Relative to neurologically asymptomatic controls, ischemic stroke patients displayed a more prevalent history of myocardial infarction and atrial fibrillation, and a higher percentage reported as taking anticoagulatory medications. Conversely, ischemic stroke patients were relatively well matched with stroke mimics in terms of all recorded cardiovascular disease risk factors, comorbidities, and medication status.

**Table 1.** Clinical and demographic characteristics.

|  | Group: |  |  | p-value: |  |  |
| --- | --- | --- | --- | --- | --- | --- |
|  | Asymptomatic<br>(A; n=20) | Stroke Mimic<br>(SM; n=20) | Ischemic Stroke<br>(IS; n=39) | Main Test | A vs. IS | SM vs. IS |
| <sup>a</sup>Age (mean ± SD) | 58.6 ± 11.1 | 58.0 ± 17.0 | 73.1 ± 13.3 | <0.001* | <0.001* | <0.001* |
| <sup>b</sup>Female n (%) | 15 (75.0) | 9 (45.0) | 25 (64.1) | 0.153 | - | - |
| <sup>a</sup>NIHSS (mean ± SD) | 0.0 ± 0.0 | 4.7 ± 4.9 | 8.6 ± 7.5 | <0.001* | <0.001* | 0.041* |
| <sup>b</sup>Family history of stroke n (%) | 11 (55.0) | 5 (25.0) | 15 (38.5) | 0.014* | 0.549 | 0.778 |
| <sup>b</sup>Hypertension n (%) | 15 (75.0) | 17 (85.0) | 32 (82.1) | 0.756 | - | - |
| <sup>b</sup>Dyslipidemia n (%) | 8 (40.0) | 13 (65.0) | 16 (41.0) | 0.199 | - | - |
| <sup>b</sup>Diabetes n (%) | 2 (10.0) | 7 (35.0) | 8 (20.5) | 0.188 | - | - |
| <sup>b</sup>Previous stroke n (%) | 1 (5.00) | 5 (25.0) | 7 (17.9) | 0.221 | - | - |
| <sup>b</sup>Atrial fibrillation n (%) | 0 (00.0) | 3 (15.0) | 13 (33.3) | 0.004* | 0.005* | 0.432 |
| <sup>b</sup>Myocardial infarction n (%) | 0 (00.0) | 6 (30.0) | 11 (28.2) | 0.014* | 0.021* | 1.000 |
| <sup>b</sup>Hypertension medication n (%) | 14 (70.0) | 16 (80.0) | 27 (69.2) | 0.723 | - | - |
| <sup>b</sup>Diabetes medication n (%) | 2 (10.0) | 6 (30.0) | 8 (20.5) | 0.309 | - | - |
| <sup>b</sup>Cholesterol medication n (%) | 6 (30.0) | 12 (60.0) | 14 (35.9) | 0.123 | - | - |
| <sup>b</sup>Anticoagulant or antiplatelet n (%) | 1 (5.00) | 12 (60.0) | 23 (59.0) | <0.001* | <0.001* | 1.000 |
| <sup>b</sup>Current smoker n (%) | 1 (5.00) | 2 (10.0) | 13 (33.3) | 0.023* | 0.043* | 0.127 |

<sup>a</sup>Means compared via one-way ANOVA with subsequent planned group-wise comparisons using Bonferroni-corrected two-sample two-tailed t-test; <sup>b</sup>Proportions compared via 2x3 Fisher's exact test with subsequent planned group-wise comparisons using Bonferroni-corrected 2x2 Fisher's exact test; NIHSS, National Institutes of Health stroke scale; SD, standard deviation

### ADAM17 activity and sCD163 levels are elevated in the peripheral blood following stroke

In order to determine whether stroke drives an increase in peripheral sCD163 production *in vivo*, peripheral venous blood was collected within 24 hours of symptom onset, and expression of ADAM17 and CD163 was assessed at both the RNA and protein level using a combination of qRT-PCR, enzyme activity assay, and ELISA. Transcriptional levels of both ADAM17 and sCD163 were significantly elevated in the whole blood of ischemic stroke patients relative to that of both control groups independently of age (Figure 1A, 1B). Protein assays supported these observations, as ischemic stroke patients displayed significantly heightened cellular ADAM17 activity in peripheral blood total hemocyte fractions (Figure 1C), as well as significantly elevated plasma levels of sCD163 (Figure 1D), once again in an age-independent fashion.

**Figure 1.**
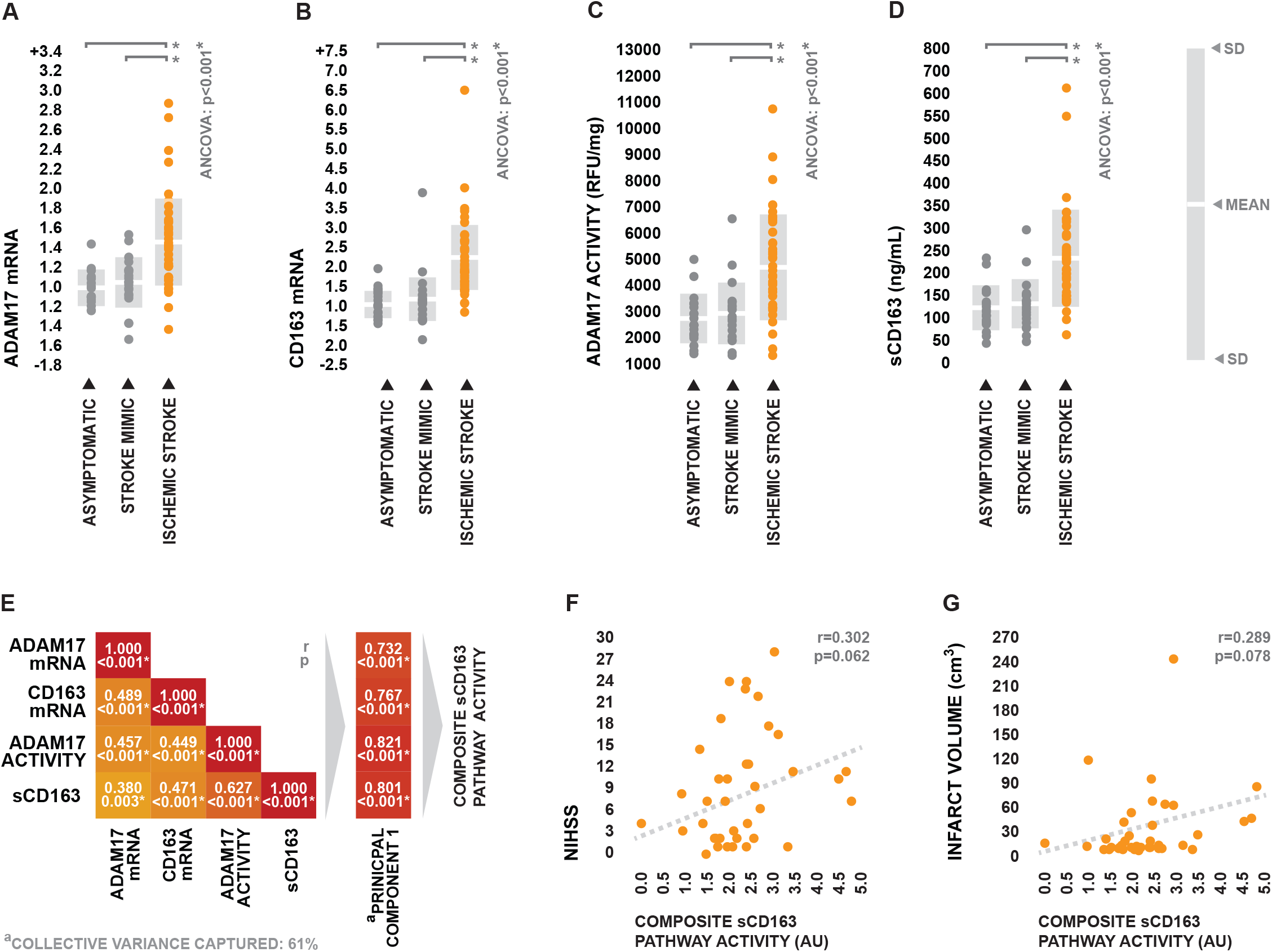
ADAM17 and CD163 expression in the peripheral blood following ischemic stroke. (A-D) Total ADAM17 mRNA expression, total CD163 mRNA expression, total cellular ADAM17 activity, and plasma sCD163 concentrations in peripheral blood biopsied from ischemic stroke patients, acute stroke mimics, and neurologically asymptomatic controls. mRNA expression values are presented as fold difference relative to asymptomatic control. Inter-group comparison of means was performed using one-way ANCOVA with age as a covariate; in the case of a significant test, subsequent planned group-wise comparisons of age-adjusted values were performed via Bonferroni-corrected two-sample two-tailed t-test. ANCOVA was performed using log-transformed data in order to meet homogeneity of variance assumptions. (E) Correlation matrix depicting the group-wide collective relationships between peripheral blood ADAM17 mRNA expression, CD163 mRNA expression, ADAM17 activity, and sCD163 levels, along with the loadings of the single principal component used to summarize sCD163 pathway activity. Strength of correlations were tested using Pearson’s r and p-values were adjusted for multiple tests via the Bonferroni correction. (F-G) Relationships between sCD163 pathway activity and injury severity as determined by NIHSS and infarct volume within the ischemic stroke group. Strength of correlations were tested using Spearman’s rho.

Expectedly, ADAM17 mRNA expression, CD163 mRNA expression, cellular TACE activity, and sCD163 levels were significantly positively correlated across the collective patient population, allowing us to summarize the peripheral activity of the sCD163 production pathway in terms of a single composite variable using principal components analysis (Figure 1E); this composite variable was positively associated with injury severity in terms of both the National Institutes of Health stroke scale (NIHSS) and infarct volume (Figure 1F, 1G). While our relatively small sample size and the variability inherent to such measures of injury status kept these associations from reaching statistical significance, these relationships tentatively inferred that the sCD163 production pathway is directly responsive to stroke pathology.

Taken as a whole, these collective observations provided associative evidence supporting our hypothesis that stroke induces a peripheral rise in sCD163 levels via increased ADAM17 activity.

### sCD163 levels are negatively associated with post-stroke lymphocyte counts

To explore whether the elevations in sCD163 levels which we observed in ischemic stroke play a role in modulation of the stroke-induced peripheral immune response *in vivo*, we assessed the relationship between circulating sCD163 levels and post-stroke leukocyte counts within the ischemic stroke group. Plasma sCD163 levels were significantly negatively associated with post-stroke absolute lymphocyte counts (Figure 2A), as well as significantly positively associated with both absolute monocyte counts and absolute neutrophil counts (Figure 2B, 2C). Collectively, these results provided *in vivo* associative evidence which supported our hypothesis that stroke-induced increases in myeloid-derived circulating sCD163 may contribute to post-stroke suppression of the adaptive immune system.

**Figure 2.**
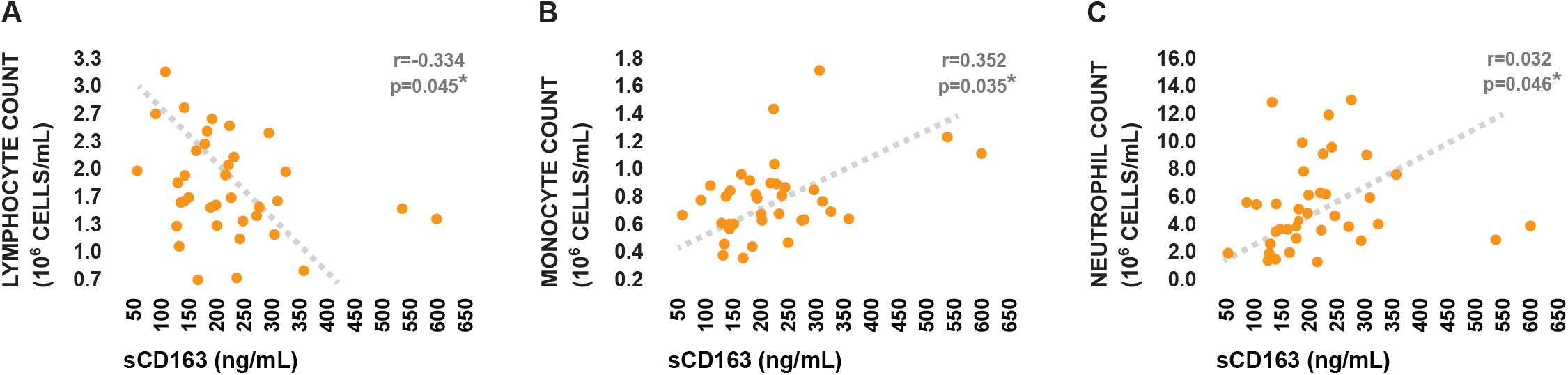
Relationship between plasma sCD163 levels and post-stroke peripheral immune status. (A-C) Relationships between plasma sCD163 levels and absolute lymphocyte counts, absolute monocyte counts, and absolute neutrophil counts in the peripheral blood of ischemic stroke patients. Strength of correlational relationships were tested via Spearman’s rho.

### Stroke-induced circulating factors trigger monocytic sCD163 production in vitro

Next, we wanted to determine the direct effects of the post-stroke peripheral inflammatory milleu on ADAM17-mediated sCD163 shedding by peripheral innate immune cell populations. To do this, primary neutrophil and monocyte cultures generated from the peripheral blood of twelve healthy donors (Supplementary Table 1) were treated with pooled serum samples derived from a subset of ten ischemic stroke patients, ten neurologically asymptomatic controls, and ten stroke mimics which were relatively well matched in terms of clinical and demographic characteristics (Table 2). Serum incubation was performed in both the presence and absence of the ADAM17 inhibitor Marimastat, and cellular ADAM17 activity and sCD163 production were assessed following three hours of treatment.

**Table 2.** Clinical and demographic characteristics of subject sub-populations used for in vitro experiments.

|  | Group: |  |  | p-value: |  |  |
| --- | --- | --- | --- | --- | --- | --- |
|  | Asymptomatic<br>(A; n=10) | Stroke Mimic<br>(SM; n=10) | Ischemic Stroke<br>(IS; n=10) | Main Test | A vs. IS | SM vs. IS |
| <sup>a</sup>Age (mean ± SD) | 57.5 ± 11.1 | 56.4 ± 17.1 | 63.5 ± 14.3 | 0.536 | - | - |
| <sup>b</sup>Female n (%) | 8 (80.0) | 5 (50.0) | 5 (50.0) | 0.348 | - | - |
| <sup>a</sup>NIHSS (mean ± SD) | 0.0 ± 0.0 | 1.6 ± 2.4 | 6.6 ± 6.9 | 0.007* | <0.001* | <0.001* |
| <sup>b</sup>Family history of stroke n (%) | 5 (50.0) | 2 (20.0) | 4 (40.0) | 0.510 | - | - |
| <sup>b</sup>Hypertension n (%) | 9 (90.0) | 8 (80.0) | 8 (80.0) | 1.000 | - | - |
| <sup>b</sup>Dyslipidemia n (%) | 5 (50.0) | 7 (70.0) | 4 (40.0) | 0.534 | - | - |
| <sup>b</sup>Diabetes n (%) | 1 (10.0) | 1 (10.0) | 1 (10.0) | 1.000 | - | - |
| <sup>b</sup>Previous stroke n (%) | 1 (10.0) | 1 (10.0) | 2 (20.0) | 1.000 | - | - |
| <sup>b</sup>Atrial fibrillation n (%) | 0 (00.0) | 2 (20.0) | 3 (30.0) | 0.321 | - | - |
| <sup>b</sup>Myocardial infarction n (%) | 0 (00.0) | 1 (10.0) | 2 (20.0) | 0.754 | - | - |
| <sup>b</sup>Hypertension medication n (%) | 9 (90.0) | 8 (80.0) | 7 (70.0) | 0.847 | - | - |
| <sup>b</sup>Diabetes medication n (%) | 1 (10.0) | 1 (10.0) | 1 (10.0) | 1.000 | - | - |
| <sup>b</sup>Cholesterol medication n (%) | 2 (20.0) | 7 (70.0) | 3 (30.0) | 0.110 | - | - |
| <sup>b</sup>Anticoagulant or antiplatelet n (%) | 1 (10.0) | 5 (50.0) | 6 (60.0) | 0.065 | - | - |
| <sup>b</sup>Current smoker n (%) | 1 (10.0) | 1 (10.0) | 2 (20.0) | 1.000 | - | - |

<sup>a</sup>Means compared via one-way ANOVA with subsequent planned group-wise comparisons using Bonferroni-corrected two-sample two-tailed t-test; <sup>b</sup>Proportions compared via 2x3 Fisher's exact test with subsequent planned group-wise comparisons using Bonferroni-corrected 2x2 Fisher's exact test; NIHSS, National Institutes of Health stroke scale; SD, standard deviation

Neutrophil cultures across all treatment conditions appeared largely phenotypically identical when visually inspected following treatment (Figure 3A). ADAM17 activity did appear to be elevated in neutrophil cultures treated with serum obtained from ischemic stroke patients relative to those treated with serum obtained from both neurologically asymptomatic controls and stroke mimics, however this increase was not statistically significant (Figure 3B). Overall, neutrophil cultures appeared to generate relatively little sCD163, although a limited number of cultures treated with ischemic stroke serum did appear to exhibit increased sCD163 production in response to treatment. This effect was not widespread however, and no statistically significant differences were observed in sCD163 production between cultures treated with ischemic stroke serum and those treated with serum from control groups (Figure 3C). A second independent experiment using pooled serum samples derived from a separate sub-set of subjects (Supplementary Table 2) yielded similar results (Supplementary Figure 1).

**Figure 3.**
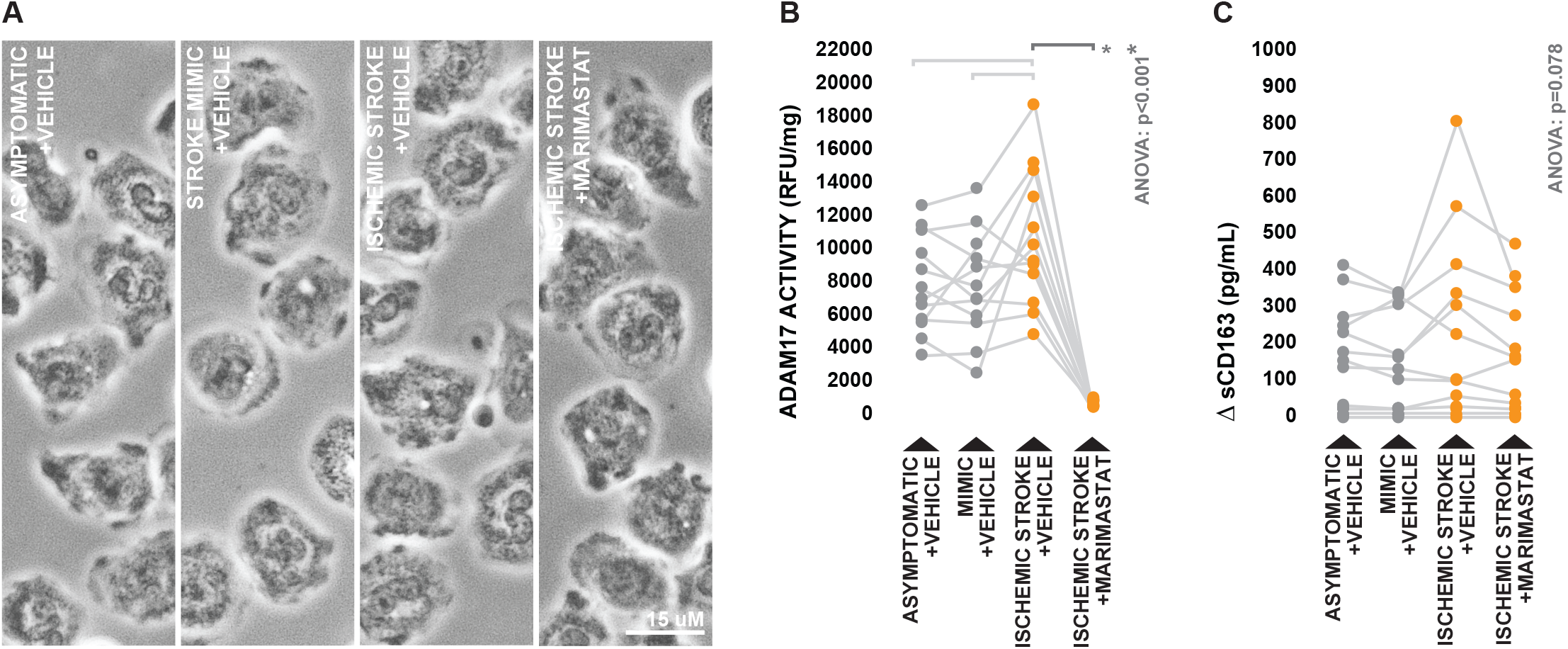
Effects of ischemic stroke serum on neutrophil ADAM17-dependant sCD163 production. (A) Morphology of healthy donor-derived neutrophils following three hours incubation with 10% serum obtained from ischemic stroke patients and control subjects, in either the presence and absence of the ADAM17 inhibitor Mari-mastat. (B) ADAM17 activity in neutrophil lysates collected following treatment. (C) Neutrophil-derived sCD163 levels in cell culture supernatants collected following treatment, presented as the difference in sCD163 levels observed between cell culture supernatants and serum-supplemented media incubated in the absence of cells. Means were compared via repeated measures one-way ANOVA; the Greenhouse-Geisser correction was applied to account for non-sphericity. In the case of a significant test, subsequent planned group-wise comparisons were performed using Bonferroni-corrected paired two-tailed t-test; planned comparisons are indicated by brackets.

Counter to what was observed in terms of neutrophil cultures, there were striking visual differences between monocyte cultures treated with serum derived from ischemic stroke patients and those treated with serum derived from control groups. Cultures treated with control serum generally displayed a mix of adherent and semi-adherent cells, however several cultures incubated with ischemic stroke serum contained large patches of cells which appeared highly adherent with prominent cytoplasmic vacuoles classically characteristic of a shift towards a macrophage phenotype (Figure 4A). ADAM17 activity was significantly elevated in monocytes treated with ischemic stroke serum relative to those treated with serum from both control groups (Figure 4B). Furthermore, supernatants recovered from cultures treated with ischemic stroke serum contained significantly higher concentrations of monocyte-derived sCD163 than those recovered from cultures treated with serum from either control group, and this effect was largely ablated by inhibition of ADAM17 (Figure 4C). Once again, similar results (Supplementary Figure 2) were observed in a second independent experiment using pooled serum samples derived from a separate sub-set of subjects (Supplementary Table 2).

**Figure 4.**
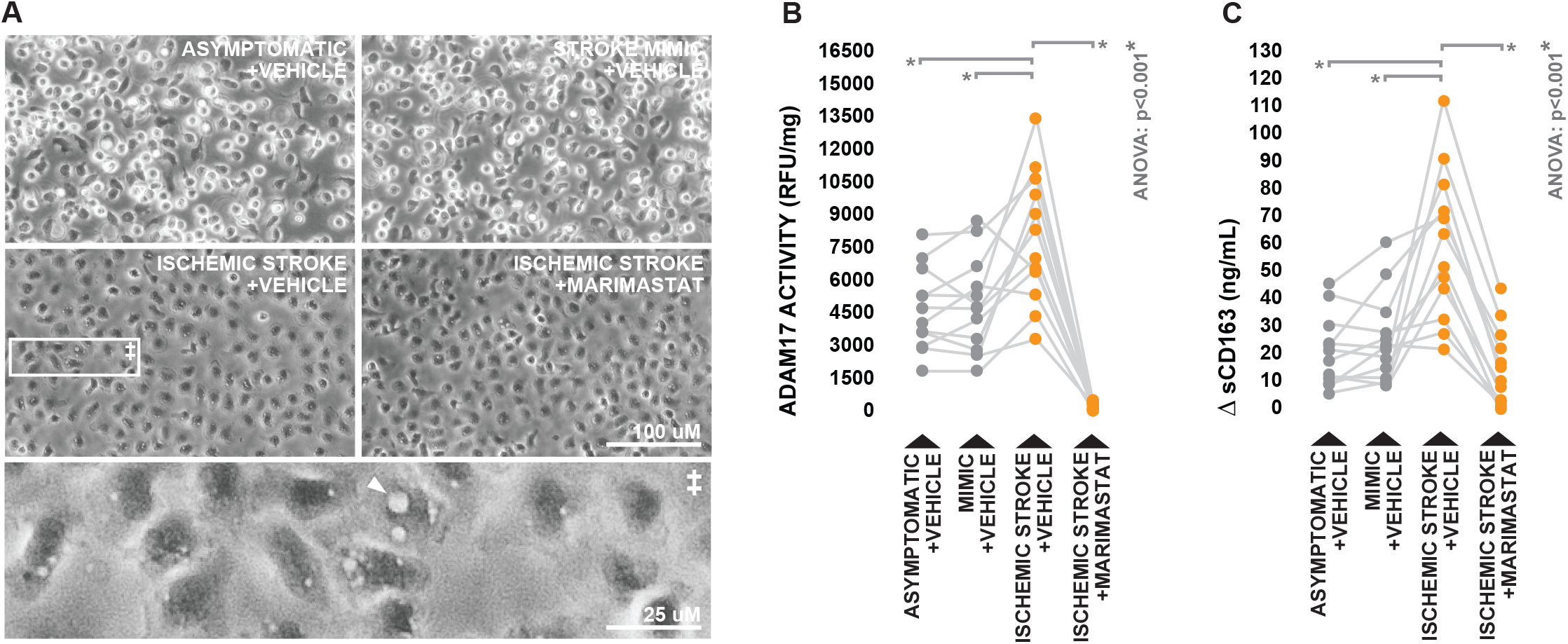
Effects of ischemic stroke serum on monocyte ADAM17-dependant sCD163 production. (A) Morphology of healthy donor-derived monocytes following three hours incubation with 20% serum obtained from ischemic stroke patients and control subjects, in either the presence and absence of the ADAM17 inhibitor Marimastat. White arrowheads indicate cytoplasmic vacuoles. (B) ADAM17 activity in monocyte lysates collected following treatment. (C) monocyte-derived sCD163 levels in cell culture supernatants collected following treatment, presented as the difference in sCD163 levels observed between cell culture supernatants and serum-supplemented media incubated in the absence of cells. Means were compared via repeated measures oneway ANOVA; the Greenhouse-Geisser correction was applied to account for non-sphericity. In the case of a significant test, subsequent planned group-wise comparisons were performed using Bonferroni-corrected paired two-tailed t-test; planned comparisons are indicated by brackets.

Collectively, these observations demonstrate that soluble factors present in peripheral circulation following ischemic stroke have the capacity to trigger ADAM17-dependant sCD163 shedding from peripheral blood monocytes, however likely not from neutrophils.

### Stroke-induced elevations in circulating sCD163 suppress lymphocyte proliferation in vitro

Lastly, we wanted to determine the direct effect of post-stroke-elevations in circulating sCD163 levels on the capacity of the peripheral blood to support lymphocyte proliferation. To do this, primary lymphocyte cultures derived from healthy donors (Supplementary Table 1) were stimulated to proliferate using phytohemagglutinin-M (PHA-M) for 72 hours in the presence of pooled serum samples generated from ischemic stroke patients, neurologically asymptomatic controls, and stroke mimics (Table 2); pooled serum samples were either unmanipulated in terms of sCD163 levels or partially depleted of sCD163 using immunoprecipitation prior to treatment (Figure 5A). BrdU incorporation was monitored over the final 24 hours of stimulation and used to assess proliferation rates.

**Figure 5.**
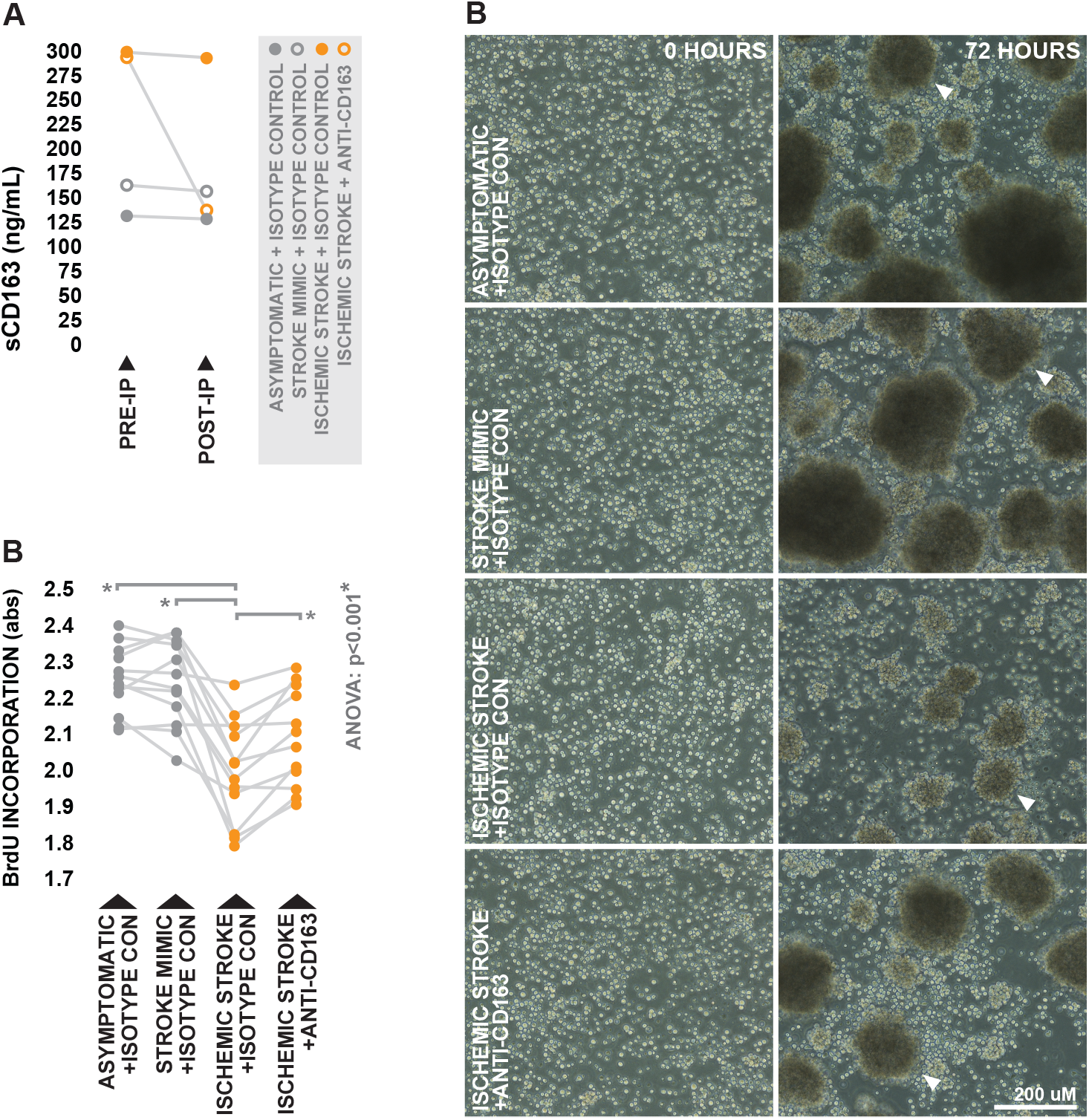
Influence of post-stroke peripheral blood sCD163 levels on the capacity to support lymphocyte proliferation. (A) Pre and post-immunoprecipitation concentrations of sCD163 in pooled serum samples obtained from ischemic stroke patients and controls, immunoprecipitated using either anti-CD163 polyclonal antibody or isotype control. (B) Morphology of healthy donor-derived lymphocytes following 72 hours of PHA-stimulated proliferation in the presence of 20% post-immunoprecipitation pooled serum. White arrowheads indicate clusters of blasting lymphocytes. (C) BrdU incorporation over the final 24 hours of treatment. Means were compared via repeated measures one-way ANOVA; the Greenhouse-Geisser correction was applied to account for non-sphericity. In the case of a significant test, subsequent planned group-wise comparisons were performed using Bonferroni-corrected paired two-tailed t-test; planned comparisons are indicated by brackets.

Consistent with prior reports,^31^ visual observations suggested decreased proliferative activity in lymphocyte cultures treated with unmanipulated ischemic stroke serum relative to those treated with unmanipulated serum from control groups, as they appeared to contain smaller and more diffuse clusters of blasting lymphocytes. However, visual indications suggested that the inhibitory effect of ischemic stroke serum was at least moderately counteracted by depletion of sCD163 (Figure 5B). In agreement with these visual observations, we observed significantly lower levels of BrdU incorporation in lymphocytes stimulated in the presence of unmanipulated ischemic stroke serum relative to those which were stimulated in the presence of unmanipulated serum derived from control groups, an effect which was partially ablated as a result of sCD163 depletion (Figure 5C). A second independent experiment using pooled serum samples derived from a separate subset of subjects (Supplementary Table 2) yielded similar results (Supplementary Figure 3). Taken together, these results novelly suggest that post-stroke elevations in circulating levels of sCD163 can have direct inhibitory effects on the proliferative capacity of lymphocytes.

## Discussion

The primary objective of this work was to determine the role of CD163 within the context of stroke immunopathology. We hypothesized that coordinate elevations in CD163 expression and ADAM17 activity induced by stroke could drive increases in circulating levels of sCD163, which could ultimately contribute to suppression of the peripheral adaptive immune system via the known inhibitory effects of sCD163 on lymphocyte activity. Our collective findings supported this hypothesis, and suggest that sCD163 plays a novel role as a factor which synergistically links stroke-induced activation of the innate immune system and suppression of the adaptive immune system. Such a mechanism provides a unique example which suggests that the innate immune system employs protective measures aimed at mitigating the risk of post-stroke autoimmune complications driven by adaptive immune system overactivation.

Observations of elevated levels of sCD163 in the peripheral blood of ischemic stroke patients, along with the fact that these elevations were negatively associated with lymphocyte counts, provided *in vivo* associative evidence supporting our hypothesis that CD163 plays a role in modulation of the adaptive immune system following stroke. Furthermore, our *in vitro* observations suggested that this stroke-driven increase in circulating sCD163 most likely originates from activated peripheral blood monocytes, as serum from ischemic stroke patients stimulated ADAM17-dependant sCD163 production in monocyte cultures generated from the peripheral blood of healthy individuals. Our *in vitro* observations further suggested that stroke-induced elevations in circulating sCD163 have the capacity to elicit direct suppressive effects on the adaptive immune system, as depletion of sCD163 from ischemic stroke patient-derived serum was able to partially rescue its capacity to support lymphocyte proliferation. Collectively, our findings infer a mechanism in which peripherally circulating factors induced by stroke trigger ADAM17-dependant CD163 shedding by monocytes, driving an increase in sCD163 levels which act to suppress peripheral lymphocyte activity; this mechanism likely serves as a means of maintaining self-tolerance as the blood brain barrier becomes disrupted and peripheral lymphoid populations become exposed to activated innate antigen presenting cells loaded with unfamiliar neural antigens (Figure 6).

**Figure 6.**
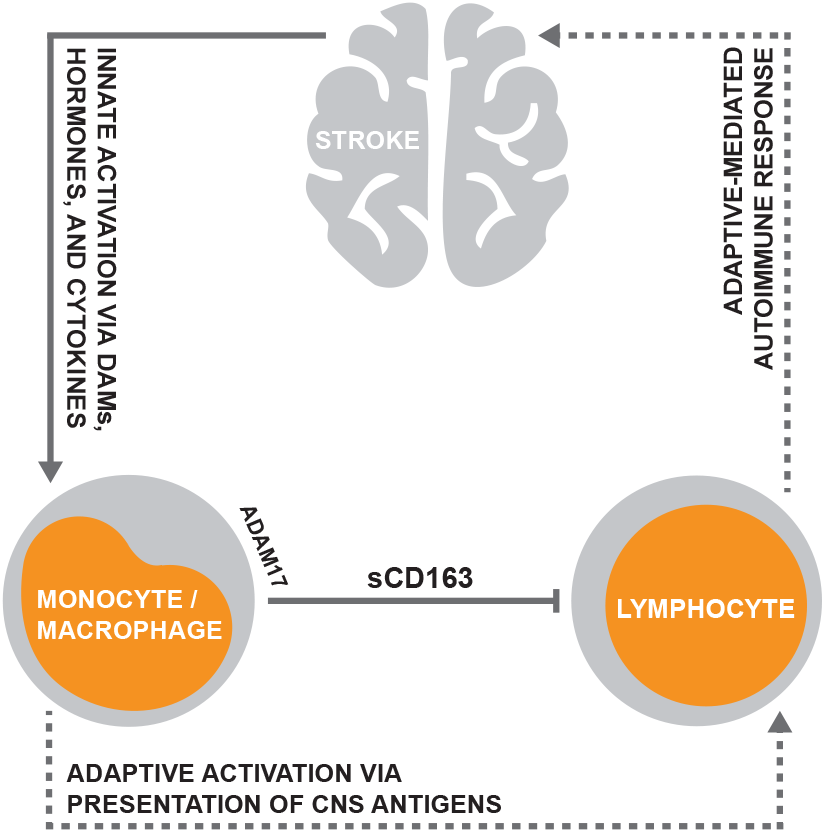
Proposed role for CD163 in modulation of the stroke-induced peripheral immune response. Stroke-driven elevations in circulating levels of damage associated molecules (DAMs), proinflammatory cytokines, and hormones coordinately trigger innate immune system activation and ADAM17-dependant CD163 shedding by monocytes, resulting in increased generation of sCD163. Heightened circulating sCD163 levels subsequently act to suppress lymphocyte activity, likely as a means of limiting the risk of a CNS-directed autoimmune response resulting from adaptive immune system activation.

To our knowledge, our results are the first to demonstrate a direct role for the innate immune system in perpetuating stroke-induced suppression of the peripheral adaptive immune system. While our results highlighted sCD163 as a means of such cross-communication, it is likely other innate-derived factors function to mediate the adaptive immune response to stroke via similar mechanisms. This notion is accentuated by the fact that ablation of sCD163 from stroke-derived serum was not adequate to fully rescue its capacity to support lymphocyte proliferation, inferring that there are a multitude of other soluble factors present in post-stroke peripheral circulation which have similar inhibitory effects. While the origins of these factors are inevitably diverse, it is likely that some arise via a similar mechanism as sCD163, as biological mechanisms are often redundant.^32^ Interesting from this regard is cluster of differentiation 14 (CD14), a membrane-bound pattern recognition receptor which is largely exclusive to the innate immune system.^33,34^ Much like CD163, CD14 can be shed via a metalloproteinase-dependent mechanism to generate a soluble protein (sCD14) which has been shown to directly interact with lymphocytes and exert suppressive effects.^35,36^ Preliminary results from our laboratory suggest that circulating levels of sCD14 are also elevated in response to stroke, and further exploration may reveal that sCD14 plays a similar role as sCD163 in modulation of the peripheral adaptive immune response.

While this study identified a clear role for sCD163 in stroke immunopathology in terms of its end-action, our experiments did not look to identify the soluble factors present in peripheral circulation following stroke which are responsible for triggering the shedding of CD163 from monocytes. It is plausible that this phenomenon is likely driven by a synergistic combination of stroke-induced damage associated molecules (DAMs), cytokines, and hormones, as members of the aforementioned families of factors are known to stimulate CD163 shedding. Notable from this standpoint is extracellular hemoglobin, a DAM which has been previously demonstrated as being acutely elevated in peripheral circulation following stroke as a result of intravascular and extravascular hemolysis.^37^ Free hemoglobin has been characterized as a potent inducer of CD163 shedding,^38^ and thus, may help drive production of sCD163 in response to stroke. Along similar lines, levels of interleukin-6, cortisol, and reactive oxygen species are all known to be elevated acutely in stroke pathology,^29,39,40^ and all have been shown to promote sCD163 production.^41,42^ Further exploration into these and other potential molecular signals which may trigger sCD163 generation within the context of stroke could not only provide a more complete picture of the mechanisms identified in this study, but a better understanding of post-stroke adaptive immune suppression as a whole, as the factors which drive post-stroke sCD163 production likely exert suppressive effects on the peripheral immune system via multiple parallel mechanisms.

While our findings are exciting, it is important to note that potentially confounding the interpretation of our results is the fact that elevations in sCD163 have been previously reported in atherosclerosis as a result of excessive intravascular macrophage activity within atherosclerotic plaques.^43^ As atherosclerosis is closely linked with stroke as one of its primary pathogenic drivers,^44^ it leaves open the possibility that our results were atherosclerosis driven, and not by the acute event of stroke itself. However, we find this scenario unlikely, as subjects in our control groups were relatively well matched with ischemic stroke patients in terms of the prevalence of cardiovascular disease risk factors associated with atherosclerosis. We suspect that sCD163 levels are chronically heightened with underlying cardiovascular pathology and become further elevated in response to the acute event of stroke itself. Supporting our notion that the elevations in sCD163 which we observed in ischemic stroke were a direct result of the stroke-induced neurological insult is the fact that it has recently been reported that levels of macrophage-derived sCD163 are elevated in the cerebrospinal fluid of pediatric traumatic brain injury patients,^45^ providing concurrent evidence that cerebral trauma has the capacity to directly trigger increased sCD163 production independent of atherosclerotic pathology.

From a broader perspective, our findings are intriguing as they demonstrate a departure from the often compartmentalized view of stroke immunopathology. While there are exceptions, research regarding stroke immunopathology has long treated the adaptive and innate immune responses as independent entities, however our results and recent results of others suggest that these responses are synergistically linked via direct mechanisms. This realization is of utmost importance with regards to the current push towards the development of novel stroke therapeutics which target the peripheral immune system. Currently, experimental immune therapies are being developed which largely aim to improve stroke outcome via one of two often discrete avenues: either by inhibiting the acute innate immune response to stroke as means of limiting excessive tissue damage and mitigating the risk of negative secondary cerebrovascular events such as edema and hemorrhagic transformation,^46–48^ or by post-acutely stimulating the peripheral adaptive immune system as a means of limiting the risk of complications induced by post-stroke infection.^49–52^ However, immunotherapeutics proposed along this line of thinking often fail to account the possible interplay between the adaptive and innate immune systems, and the potential repercussions that could stem from modulating one response without consideration for the other.

A perfect example in this regard is the fact that ADAM17 inhibitors are often cited as a potential stroke therapeutic under the premise that they could ultimately limit excessive innate-driven inflammation and its associated complications via a reduction in TNFα production.^53–56^ However, our results clearly demonstrate that inhibition of ADAM17 within the innate immune system could directly interfere with mechanisms which act to suppress the adaptive immune system, an unintended effect which could put patients at elevated risk for the development of long-term autoimmune complications associated with loss of CNS self-tolerance. Such an example highlights the necessity for a global view regarding the peripheral immune system in terms of the development of novel stroke immunotherapeutics. Unfortunately, therapeutic design from such a perspective is currently unrealistic, as the mechanisms which drive the stroke-induced peripheral immune response, particularly that of the adaptive immune system, are still largely unknown. This current gap in knowledge may underlie the reality that stroke immunotherapeutics have thus-far been largely unsuccessful.^57^ Further future work which aims to characterize the peripheral immune response to stroke and its underlying mechanisms from a panoptic systems biology approach, preferably within human-centric models, would likely generate the foundational basis for the development of stroke immunotherapeutics with higher chances for clinical success.

Collectively, our results demonstrate a novel role for CD163 as factor which mechanistically couples stroke induced-activation of the peripheral innate immune system and suppression of the peripheral adaptive immune system. Our results provide a unique example which suggests that the innate immune system employs protective mechanisms aimed at mitigating the risk of post-stroke autoimmune complications driven by adaptive immune system overactivation. Further work which looks to characterize similar pathways of cross-communication between the innate and adaptive immune systems within the context of stroke would likely provide a better understanding of stroke pathology as a whole, and allow for more informed development of stroke-targeted immunotherapeutics.

## Experimental procedures

### Patients

Acute ischemic stroke patients, acute stroke mimics, and neurologically asymptomatic controls were recruited at Ruby Memorial Hospital, Morgantown, WV. Ischemic stroke patients displayed definitive radiographic evidence of vascular ischemic pathology on magnetic resonance imaging (MRI) or computed tomography (CT) according to the established criteria for diagnosis of acute ischemic cerebrovascular syndrome (AICS)^58^, and diagnoses were confirmed by an experienced neurologist. Patients admitted to the emergency department as suspected strokes based on the overt presentation of stroke-like symptoms, but receiving a definitive negative diagnosis for stroke upon neuroradiological imaging and clinical evaluation were identified as acute stroke mimics.^58^ Discharge diagnoses of stroke mimics included cases of seizures, complex migraines, hypertensive encephalopathy, and other non-stroke conditions which induce neurological symptoms. Prospective subjects were excluded if they received a non-definitive diagnosis, displayed evidence of hemorrhagic pathology, presented with cancer as a co-pathology, reported a prior hospitalization within 90 days, or were under 18 years of age. Blood was sampled within 24 hours of onset, as determined by the time the patient was last known to be free of stroke-like symptoms. In the case of patients who received thrombolytic therapy, blood samples were collected prior to the administration of recombinant tissue plasminogen activator (rtPA). Injury severity was determined according to the NIHSS at the time of blood draw. Control subjects were deemed neurologically normal by a trained neurologist at the time of enrolment. Demographic information was collected from either the subject or significant other by a trained clinician. Procedures were approved by the institutional review boards of West Virginia University and Ruby Memorial Hospital (IRB protocol 1410450461R001), and all experiments were performed in accordance with relevant guidelines and regulations. Written informed consent was obtained from all subjects or their authorized representatives prior to study procedures.

### Neuroradiological imaging

Neuroradiological imaging was performed using either MRI or CT within 24 hours of symptom onset. CT imaging was performed on either a Toshiba Aquilion 64 or Toshiba Aquilion 320 scanner (5 mm slices acquired at 120 KVp, 250 mA). MRI imaging was performed using either a Siemens Aera or Siemens Verio scanner (5 mm slices, 5 mm inter-slice gap, acquired via diffusion-weighted echoplanar imaging with a B-factor of 1000 s/mm^2^). The BrainLAB iPlan Neuroradiology software package (BrainLAB, Westchester, Ill) was used to calculate infarct volume via manual tracing as previously described,^59^ and all infarct volume calculations were verified by an experienced neurologist.

### Blood collection

Parallel peripheral venous whole blood samples were collected from subjects via PAXgene RNA tubes (Qiagen, Valencia, CA), K2 EDTA Vacutainers (Becton Dickenson, Franklin Lakes, NJ), and serum separator tubes (Becton Dickenson). PAXgene RNA tubes were frozen immediately and stored at −80°C until RNA extraction. K2 EDTA tubes were stored at room temperature until white blood cell differential (less than 30 minutes), plasma isolation (less than 30 minutes), or leukocyte isolation (performed immediately). Serum separator tubes were stored at room temperature until serum isolation (less than 30 minutes).

### Isolation of serum and plasma

For plasma isolation, EDTA-treated blood was spun at 2,000*g for 10 minutes to sediment hemocytes. Plasma was collected for ELISA, and the total hemocyte fraction was retained for subsequent protein extraction and ADAM17 activity assay. Plasma was additionally spun at 10,000*g for 10 minutes to remove any residual blood cells or debris. For serum isolation, serum separator tubes were incubated at room temperature for a minimum of 15 minutes to allow for coagulation, and subsequently spun at 2,000*g for 10 minutes to sediment the resulting thrombus. Resultant samples were stored at −80°C until analysis.

### RNA extraction and quantitative reverse transcription PCR

Whole blood RNA was extracted from PAXgene tubes via the PreAnalytiX PAXgene blood RNA kit (Qiagen) and automated using the QIAcube system (Qiagen). RNA was isolated from leukocyte lysates via the RNeasy Micro kit (Qiagen). Quantity and purity of isolated RNA was determined via spectrophotometry (NanoDrop, Thermo Scientific, Waltham, MA).

cDNA was generated from purified RNA using the Applied Biosystems high capacity reverse transcription kit. For qPCR, target sequences were amplified from 10 ng of cDNA input using sequence specific primers (Supplementary Table 3) and detected via SYBR green (PowerSYBR, Thermo-Fisher) on the RotorGeneQ (Qiagen). Raw amplification plots were background corrected and CT values were generated via the RotorGeneQ software package. All reactions were performed in triplicate. Transcripts of *B2M, PPIB*, and *ACTB* were amplified as references and normalization was performed using the NORMA-Gene data-driven normalization algorithm.^60^

### White blood cell differential

Complete blood count was obtained from EDTA-treated blood via combined optical flow cytometry and cellular impedance using an automated clinical hematology analyzer (Cell-Dyn, Abbott Diagnostics, Santa Clara, CA).

### Isolation of primary leukocytes for cell culture

Starting with 30 mLs of healthy donor-derived EDTA-treated blood, peripheral blood mononuclear cells (PBMCs) were separated from polymorphonuclear cells (PMNs) and red blood cells (RBCs) via centrifugation over a polysaccharide-sodium diatrizoate density gradient (Lymphoprep, 1.077 g/mL, StemCell Technologies) for 30 minutes at 400*g. Resultant interphase PBMC fractions were collected, resuspended in RPMI 1640 (Life Technologies, Grand Island, NY) containing 10% autologous serum, separated into two aliquots, and set aside at room temperature. Pelleted PMN/RBCs were immediately placed on ice for PMN enrichment via RBC lysis induced by two 5 minute incubations with a 1:10 volume of ice cold ammonium chloride (ACK) buffer. The resultant PMNs were rinsed in ice cold phosphate buffered saline (PBS), counted via automated cytometer (Cellmeter X1, Nexcelom Bioscience, Lawrence, MA), and immediately plated for experiments.

Following PNM isolation, monocytes were further enriched from one aliquot of PBMCs via a second round of density gradient centrifugation as described by Menck et al.^61^ Briefly, PBMC suspensions were layered over an iso-osmotic 46% Percoll (1.131 g/mL, Sigma Aldrich) solution prepared in RPMI 1640 containing 10% autologous serum and centrifuged at 500*g for 30 minutes. Monocytes were collected from the resultant interphase, rinsed in PBS, counted, and immediately plated for experiments.

Following PMN and monocyte isolation, the remaining aliquot of PBMCs was rinsed in PBS and seeded in T75 flasks for lymphocyte expansion. Lymphocytes were expanded for 72 hours under standard mammalian cell culture conditions in a growth media comprised of RPMI 1640 containing 2% PHA-M (Gibco, Grand Island, NY), 2 mM L-Alanyl-Glutamine (GlutaMAX, Gibco), 25 mM HEPES (Gibco), 50 uM BME (Gibco), and 1% antibiotic-antimycotic (Gibco), supplemented with 10% autologous serum. Following expansion, actively proliferating lymphocytes were rinsed in PBS, counted, and plated for experiments.

### Neutrophil culture

Neutrophils from healthy donors were seeded in 12-well plates at a density of 7.5*10^5^ cells per well in 250 uL of GlutaMax-supplemented RPMI 1640 containing 10% patient serum along with either Marimastat (10 uM, Sigma Aldrich, St-Louis, MO) or vehicle (DMSO, Fisher Scientific). Following three hours of incubation under standard mammalian culture conditions, cells were lysed in NP40 lysis buffer (Life Technologies) and cellular ADAM17 activity was assessed via enzyme activity assay. Cell culture supernatants were collected and concentrations of sCD163 were quantified via ELISA; levels of neutrophil-derived sCD163 were determined by subtracting the concentration of sCD163 in media incubated without cells from the concentrations of sCD163 observed in cell culture supernatants.

### Monocyte culture

Monocytes from healthy donors were seeded in 12-well plates at a density of 1*10^6^ cells per well in 250 uL of GlutaMax-supplemented RPMI 1640 containing 20% patient serum along with either Marimastat (10 uM) or vehicle (DMSO). Following three hours of incubation under standard mammalian culture conditions, cells were lysed in NP40 lysis buffer and cellular ADAM17 activity was assessed via enzyme activity assay. Cell culture supernatants were collected and concentrations of sCD163 were quantified via ELISA; levels of monocytes-derived sCD163 were determined by subtracting the concentration of sCD163 in media incubated without cells from the concentrations of sCD163 observed in cell culture supernatants.

### sCD163 depletion from human serum

sCD163 was depleted from pooled serum samples via immunoprecipitation using biotinylated goat polyclonal antibody raised against the extracellular region of CD163 (BAF1607, R&D Systems, Minneapolis, MN). Biotinylated normal goat IgG (BAF108, R&D Systems) was used as an isotype immunoprecipitation control. Immunoglobulins were conjugated to polymer-coated superparamagnetic beads (Dynabeads Streptavidin T1, Thermo Fisher, Waltham MA) at a ratio of 10 ug of immunoglobulins per 1 mg of beads. For immunoprecipitation, 5 mL of pooled serum was precleared with normal IgG conjugated beads, and then incubated with 0.5 mg of either normal IgG conjugated beads or anti-CD163 conjugated beads for 30 minutes at 4°C. Beads were subsequently separated from serum samples via multiple rounds of magnetic separation. Immunoprecipitation was performed under aseptic conditions and serum was filtered at 14 microns following immunoprecipitation to remove potential microbial contaminants prior to cell culture. Depletion of sCD163 was subsequently confirmed via ELISA.

### Lymphocyte proliferation assay

Expanded lymphocytes from healthy donors were seeded in 48 well plates at 5*10^4^ cell per well in 200 uL growth media supplemented 20% pooled patient serum. Following 48 hours of culture, BrdU (Roche Life Sciences, Indianapolis, IN) was added to cultures at a final concentration of 20 uM. After allowing for 24 hours for BrdU incorporation, lymphocytes were adhered via centrifugation and fixed for BrdU quantification. Incorporated BrdU was quantified via colorimetric ELISA (Roche Life Sciences) per manufacture instructions.

### sCD163 ELISA

sCD163 was measured in plasma samples and cell culture supernatants using a commercially available colorimetric ELISA assay (RAB0082, Sigma-Aldrich). Plasma samples were diluted 1:50, neutrophil cell culture supernatants were diluted 1:5, and monocyte cell culture supernatants were diluted 1:10 prior to analysis. Absorbance readings were obtained using the Synergy HT multi-mode microplate reader (BioTek, Winooski VT). Due to the potential confound of hemolysis on concentrations of leukocyte derived analytes, plasma samples were screened for hemolysis prior to ELISA. Plasma absorbance was measured at 385 and 414 nm via spectrophotometry (NanoDrop, Thermo Scientific, Waltham, MA) and used to calculate a hemolysis score as described by Appierto *et al*.^62^ Plasma samples with detectable hemolysis were excluded from analysis.

### ADAM17 activity assay

Total hemocyte fractions were thawed, mixed with NP-40 lysis buffer (Thermo Fisher) at a 1 to 1 ratio, and depleted of hemoglobin via the HemogloBind hemoglobin removal kit (Biotech Support Group, Monmouth Junction, NJ) as described by Park et al.^63^ Protein concentrations of hemoglobin-depleted total hemocyte lysates and cell culture lysates were determined via DC-protein assay (Bio-Rad, Hercules, CA). ADAM17 activity was measured in 200 ug of total hemocyte lysate, or 10 ug of cell culture lysate, via the InnoZyme fluorometric TACE activity kit (EMD Millipore, Temecula, CA) per manufacture instructions. Fluorometric readings were obtained using the Synergy HT multi-mode microplate reader (BioTek). Due to the reversible nature of ADAM17 inhibition by Marimastat, experimental concentrations of drug were maintained in assay buffers when testing samples from inhibitor-treated conditions.

### Phase Contrast Microscopy

Phase contrast images of primary leukocyte cultures were obtained from an AxioObserver inverted microscope (Ziess, Thornwood, NY) equipped with a AxioCam mR5 digital camera (Ziess) using the AxioVision imaging software suite (Ziess). Scale bars were generated via imaging of a 0.07-1.50 mm scale calibration slide (Motic, Richmond, BC, CA).

### Statistical analysis

Statistics were performed using SPSS (IBM, Chicago, Ill) in combination with R 2.14 (R project for statistical computing)^64^ via the SPSS R integration plug-in. Fisher’s exact test was used for comparison of dichotomous variables. Student t-test, one-way ANOVA, one-way ANCOVA, or repeated measures one-way ANOVA was used for comparison of continuous variables where appropriate. In some instances of ANCOVA and one-way ANOVA, data were log-transformed prior to analysis in order to meet normality and homogeneity of variance assumptions, which were evaluated via the Shapiro-Wilk test and Levene’s test. In some instances of repeated measures one-way ANOVA, values were adjusted via the Greenhouse-Geisser correction to account for nonsphericity, which was assessed using Mauchly’s test. Either Spearman’s rho or Pearson’s r was used to assess the strength of correlational relationships where appropriate based on level of homoscedasticity. Principle components analysis was used for dimensionality reduction; prior to analysis, data were tested for sphericity and sampling adequacy via Bartlett’s test and the Kaiser-Mayer-Olkin index. The level of significance was established at 0.05 for all statistical testing. In the cases of multiple comparisons, p-values were adjusted using Bonferroni correction. Parameters of all statistical tests performed are outlined in detail within the figure legends.

## Acknowledgments

The authors would foremost like to thank the subjects and their families, as this work was truly made possible by their selfless contribution. We would further like to thank the stroke team at Ruby Memorial Hospital for their support. In addition, we would like to thank Dr. Daniel Laskowitz (Duke University), Dr. Jason Huber (West Virginia University), Dr. Lori Hazlehurst (West Virginia University), and Dr. Gordon Meares (West Virginia University) for critical review of the manuscript. Finally, we would like to thank Dr. Ashley Petrone (West Virginia University) and Dr. Asano Shinichi (West Virginia University) for general laboratory assistance. Work was funded via a Robert Wood Johnson Foundation Nurse Faculty Scholar award to TLB (70319) and a National Institutes of Health CoBRE sub-award to TLB (P20 GM109098).

## Author Contributions

Work was conceptualized by GCO. Clinical sample collection and recruitment of human subjects was performed by NL-W and CST, and overseen by PDC and TLB. Analysis of neuroradiological imaging was performed by YK and ART. Molecular experiments were designed and performed by GCO. Data were analyzed by GCO. Manuscript was written by GCO with contributions from NL-W, CST, YK, ART, PDC, and TLB.

